# Recurrent Viral Capture of Cellular Phosphodiesterases that Antagonize OAS-RNase L

**DOI:** 10.1101/2023.05.12.540623

**Authors:** Stephen A. Goldstein, Nels C. Elde

**Author notes:** Co-corresponding authors –.

## Abstract

Phosphodiesterases (PDEs) encoded by viruses are putatively acquired by horizontal transfer of cellular PDE ancestor genes. Viral PDEs inhibit the OAS-RNase L antiviral pathway, a key effector component of the innate immune response. Although the function of these proteins is well-characterized, the origins of these gene acquisitions is less clear. Phylogenetic analysis revealed at least five independent PDE acquisition events by ancestral viruses. We found evidence that PDE-encoding genes were horizontally transferred between coronavirus genera. Three clades of viruses within *Nidovirales*: merbecoviruses (MERS-CoV), embecoviruses (OC43), and toroviruses encode independently acquired PDEs, and a clade of rodent alphacoronaviruses acquired an embecovirus PDE via recent horizontal transfer. Among rotaviruses, the PDE of Rotavirus A was acquired independently from Rotavirus B and G PDEs, which share a common ancestor. Conserved motif analysis suggests a link between all viral PDEs and a similar ancestor among the mammalian AKAP7 proteins despite low levels of sequence conservation. Additionally, we used ancestral sequence reconstruction and structural modeling to reveal that sequence and structural divergence are not well-correlated among these proteins. Specifically, merbecovirus PDEs are as structurally divergent from the ancestral protein and the solved structure of human AKAP7 PDE as they are from each other. In contrast, comparisons of Rotavirus B and G PDEs reveal virtually unchanged structures despite evidence for loss of function in one, suggesting impactful changes that lie outside conserved catalytic sites. These findings highlight the complex and volatile evolutionary history of viral PDEs and provide a framework to facilitate future studies.

## Introduction

Horizontal gene transfer (HGT) is a major force in ancient and ongoing evolution among diverse viruses and hosts. Within the virosphere, the capture and re-purposing of host genes has shaped evolutionary history spanning from the origins of viruses themselves (1) to more specialized interactions with host immune pathways. Viruses in distinct unrelated orders, Nidoviruses and Rotaviruses, have independently captured host mRNAs encoding eukaryotic LigT-like 2H-phosphoesterases, specifically 2’-5’ phosphodiesterases (PDEs) with similarity to eukaryotic protein family members (2). PDEs encoded by betacoronaviruses, toroviruses, and group A rotaviruses function as potent antagonists of the OAS-RNase L pathway with important roles *in vitro* and *in vivo* promoting viral replication and pathogenesis (3–8).

In contrast to the extensive research on PDE function, the origin and evolutionary history of viral PDEs remain obscure. Based on structural similarity the PDEs encoded by mouse hepatitis virus (9) and group A rotaviruses (10,11) have been linked to the PDE domain of mammalian AKAP7 proteins. Previously predicted structures for PDEs encoded by MERS-CoV (6), toroviruses (5) and group B and G rotaviruses (11) reveal an AKAP7 PDE-like fold and retain catalytic histidine (HxS/T) motifs.

The interface between viral PDEs and the OAS-RNase L pathway is an attractive opportunity to study evolutionary pressures related to the acquisition, fixation, and evolution of host-derived viral genes. The restriction of rotavirus (8) and coronavirus (3,7,12) replication in the absence of an active PDE along with the presence of these genes across diverse viruses provide strong evidence for potential fitness advantages conferred by PDE acquisition.

The capacity for viruses to accommodate increased genome size is a key determinant of their ability to evolve via HGT. Considered broadly, the genomic architecture of viral PDEs is of two types. PDEs encoded by rodent alphacoronaviruses (13), embecoviruses (5), and merbecoviruses (6) are ∼240-300 amino acids long, translated off of a discrete viral mRNA, presumably exist in the cell as autonomous proteins, and exhibit high amino acid sequence divergence relative to conserved regions of the replicase gene. However, PDEs may also act as functional protein sub-domains of as few as ∼115 amino acids. The rotavirus A PDE consists of 142 amino acids, rotavirus B 115 amino acids, and rotavirus G 116 amino acids with all organized as the C-terminal subdomain of the >750 amino acid VP3 protein while the torovirus PDE is a 141 amino acid C-terminal domain of the >4,000 amino acid polyprotein 1a, though whether it is proteolytically processed into an autonomous protein is unclear.

In this study we identified several striking features at play in the evolution of viral PDEs. We used a combination of phylogenetics, conserved motif analysis, and AlphaFold-based structural predictions to characterize the origin of viral PDEs, their diversification following horizontal transfer, and the relationship of divergent viral PDEs to each other and mammalian AKAP7. Despite similarity in function and predicted structure with each other and mammalian AKAP7, we exclude the possibility of a single horizontal transfer event giving rise to PDEs and instead propose a more complex and volatile process of diversification. These findings advance our understanding of the history of PDEs as diverse and potent immune antagonists and provide a detailed characterization of how host-derived viral genes originate and evolve.

## Results

### Viral PDEs derive from several AKAP-like horizontal transfer events

We aligned 173 viral PDE amino acid sequences (18 Torovirus, 10 Alphacoronavirus, 55 Embecovirus, 27 Merbecovirus, 8 Rotavirus G, 20 Rotavirus B, 35 Rotavirus A) using MAFFT (version 7), which due to high sequence divergence involved intensive manual refinement (see Methods for details). Next, we constructed an unrooted maximum-likelihood phylogenetic tree (**Figure 2A**). The PDE tree contains five major branches reflective of five putative acquisition events: rotavirus A (RVA) VP3, the rotavirus B (RVB) and rotavirus G (RVG) VP3 single common ancestor, toroviruses, embecoviruses, and merbecoviruses. The arrangement of some branches suffered low bootstrap support likely owing to sequence divergence, but the topology was sufficiently robust to reveal major events in viral PDE evolution.

**Figure 1.**
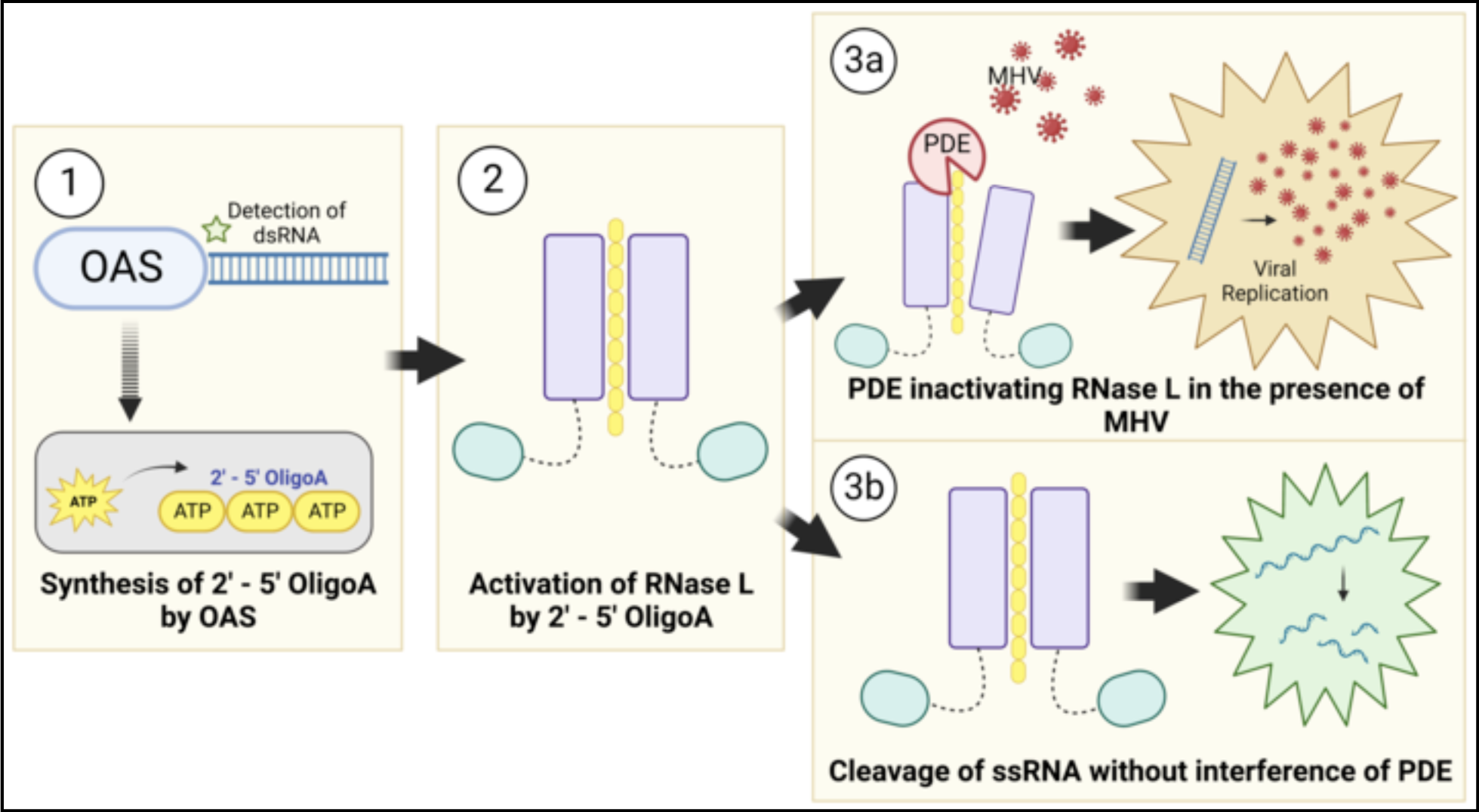
Viral PDE antagonism of OAS-RNase L. (1) OAS proteins sense viral dsRNA upon infection and synthesize 2’-5’ oligoadenylate (2-5A) by linking ATP molecules via the 2’ and 5’ carbons. (2) 2-5A binds RNase L monomers, inducing homodimerization and activation (3A) Viral PDEs prevent RNase L activation by cleavage of 2-5A, facilitating viral replication (3B) In the absence of a functional PDE RNase L is activated and cleaves both viral and cellular ssRNA, restriction viral replication, inducing translational arrest, and ultimately resulting in cell death.

**Figure 2.**
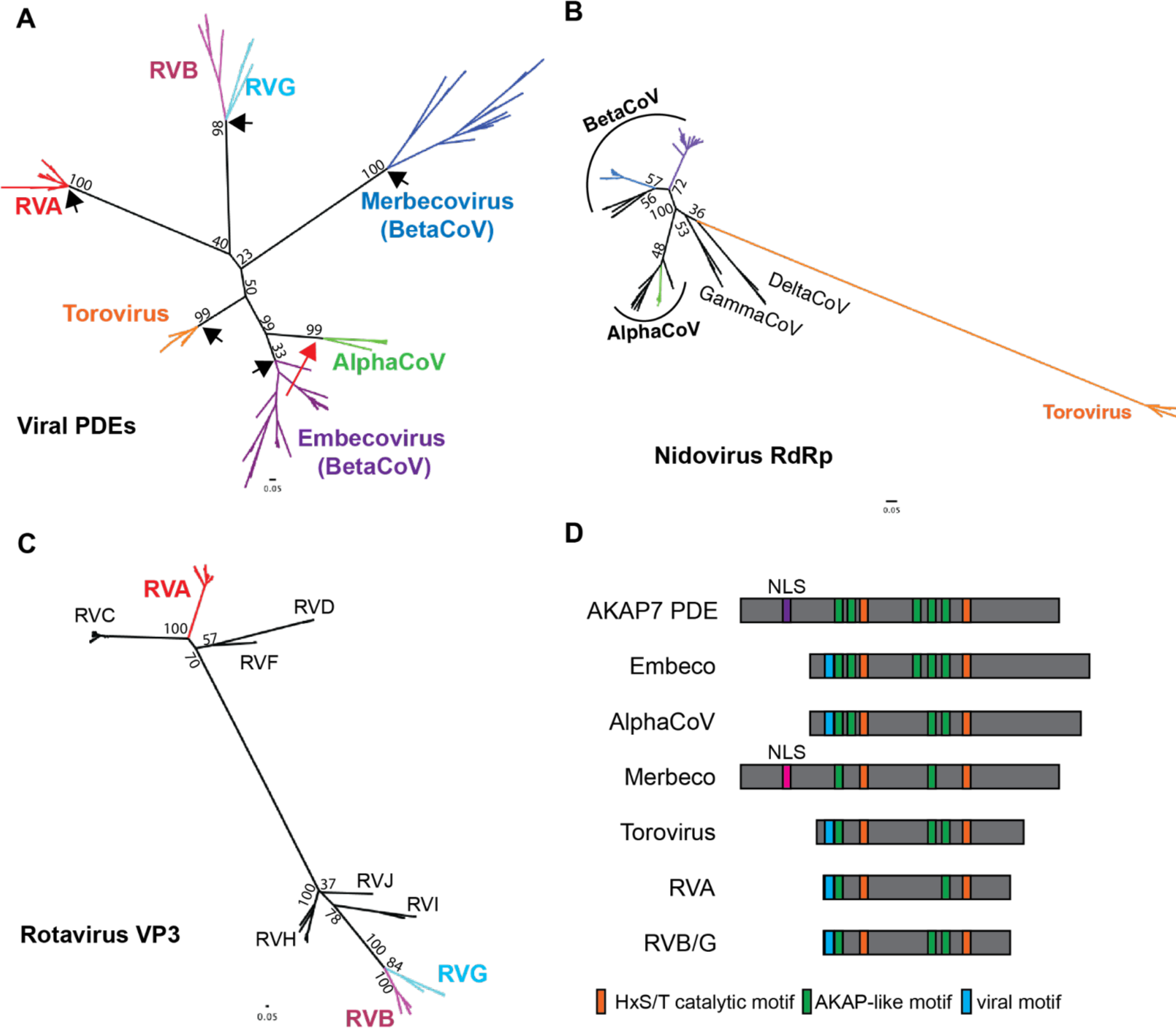
Phylogenetic and sequence analysis of viral PDEs reveals multiple independent acquisitions and conservation of ancestral motifs. A) Unrooted maximum-likelihood (ML) Tree of viral phosphodiesterase amino acid sequences showing five PDE acquisitions from a cellular ancestor (black arrows) and one acquisition via recombination (red arrow) B) Unrooted ML tree of coronavirus and torovirus full length RdRp amino acid sequences C) Unrooted ML tree of rotavirus VP3 amino acid sequences, with the C-terminal domain PDEs trimmed from the RVA, RVB, and RVG sequences D) Schematic showing the conservation or absence of AKAP-like motifs in viral phosphodiesterases.

We found evidence for three independent acquisitions of PDEs in the order *Nidovirales*. The *Nidovirales* PDE phylogeny is incongruent with the RdRp phylogeny (**Figure 2B**) and the PDEs are not syntenic between virus classes, with one exception. Consequently, we found no support for a recent common ancestor of all nidovirus PDEs. In contrast to the three acquisitions of an ancestral cellular PDE, the rodent alphacoronavirus PDE and the Embecovirus PDE are syntenic and have high sequence identity. This synteny suggests minimal homology-guided recombination played a role in the transfer of a PDE gene from one to the other. Embecovirus PDEs exhibit higher within-clade diversity, suggesting an older origin and providing support for the hypothesis that alphacoronavirus PDEs were acquired via recombination with embecoviruses (13), in contrast to the idea that they were acquired independently (**Figure 2A**) (14). Given the overlapping ecological niches of rodent betacoronaviruses and alphacoronaviruses, we cannot exclude the possibility that ancestral viruses in each clade independently acquired a similar cellular PDE. Given the greater intra-clade divergence among embecovirus which supports an older origin, however, it is unlikely independently acquired PDEs would retain such high identity.

The Rotavirus phylogeny supports with high confidence two independent PDE captures - one on the RVA VP3 branch and one into an RVB/RVG VP3 common ancestor. We constructed a maximum-likelihood tree of 137 rotavirus VP3 genes representing all known rotavirus genotypes (RVA, RVB, RVC, RVG, RVH, RVI, RVJ) with the PDE domain trimmed from the RVA, RVB, and RVG VP3s (**Figure 2C**). The VP3 phylogeny matches the PDE phylogeny where RVA is extremely distant from RVB and RVG, which derive from a recent common ancestor. This supports a parsimonious scenario in which that ancestor acquired a PDE which diversified at the same rate as the rest of the gene. Surprisingly, despite the synteny of the RVA and RVB/RVG PDEs, their origins are clearly independent of each other.

Previous work identified mammalian AKAP7 central domains (CD) as structural and functional homologs of viral PDEs (15,16). However, low sequence conservation leaves the evolutionary link between the viral and cellular PDEs uncertain. While generating the viral PDE phylogeny (**Figure 2A**) we identified several AKAP-like motifs largely conserved in the viral PDEs (**Figure 2D**) that proved critical to anchoring the low-identity alignment and conclusively link the viral PDEs to a cellular AKAP ancestor, as no other eukaryotic PDEs share these motifs. One short motif (blue in **Figure 2D**) is shared among all viral PDEs with the exception of merbecoviruses but is not present in any AKAP7. Though convergent evolution is possible, it seems more likely that this is a remnant of the collapse of the AKAP N-terminus that removed the nuclear localization sequence but maintained some function given its conservation among viral PDEs.

Finally, a PDE encoded by Orf4 of a shrew alphacoronavirus was recently reported (14), raising the possibility of an additional independent PDE acquisition, or virus-to-virus horizontal transfer. Its phylogenetic positioning remains uncertain on the end of a long branch with very low support due to sparse sampling (**Figure S1A**). Several of the key motifs, including the second catalytic motif, are degraded, indicating that this protein is almost certainly non-functional (**Figure S1B**). This observation further highlights the prevalence of PDE acquisitions by horizontal gene transfer.

### Ancestral Sequence Reconstruction of Nidovirus PDEs

Our phylogenetic analysis revealed clear distinctions between Nidovirus PDE clades, supporting independent acquisition of PDEs by Merbecoviruses, Embecoviruses, and Toroviruses. Nevertheless, confidence is clouded by the high sequence divergence among PDEs and the shared, recombination-driven history of frequent recombination between Embecovirus and Alphacoronavirus PDEs. Consequently, we performed additional analysis combining ancestral sequence reconstruction with AlphaFold2 structural modeling to further investigate relationships among viral PDEs (17). We initially focused on Embecovirus NS2 and Merbecovirus NS4b proteins given the higher degree of within-clade sequence diversity among these proteins, which likely indicates a combination of more thorough sampling and an older acquisition. We aligned 55 Embecovirus NS2 sequences with the *Hs*AKAP7 PDE domain. We then constructed a maximum-likelihood tree of Embecovirus NS2 protein sequence with an *Hs*AKAP7 outgroup rooting and used this tree as the basis for ancestral reconstruction in FastML (**Figure 3A**). We aligned the *Hs*AKAP7 and ancestral Embecovirus NS2 sequences from major nodes and calculated amino acid identity. The deepest ancestral node exhibited substantially higher amino acid identity to *Hs*AKAP7 than shallower nodes or observed extant sequences.

**Figure 3.**
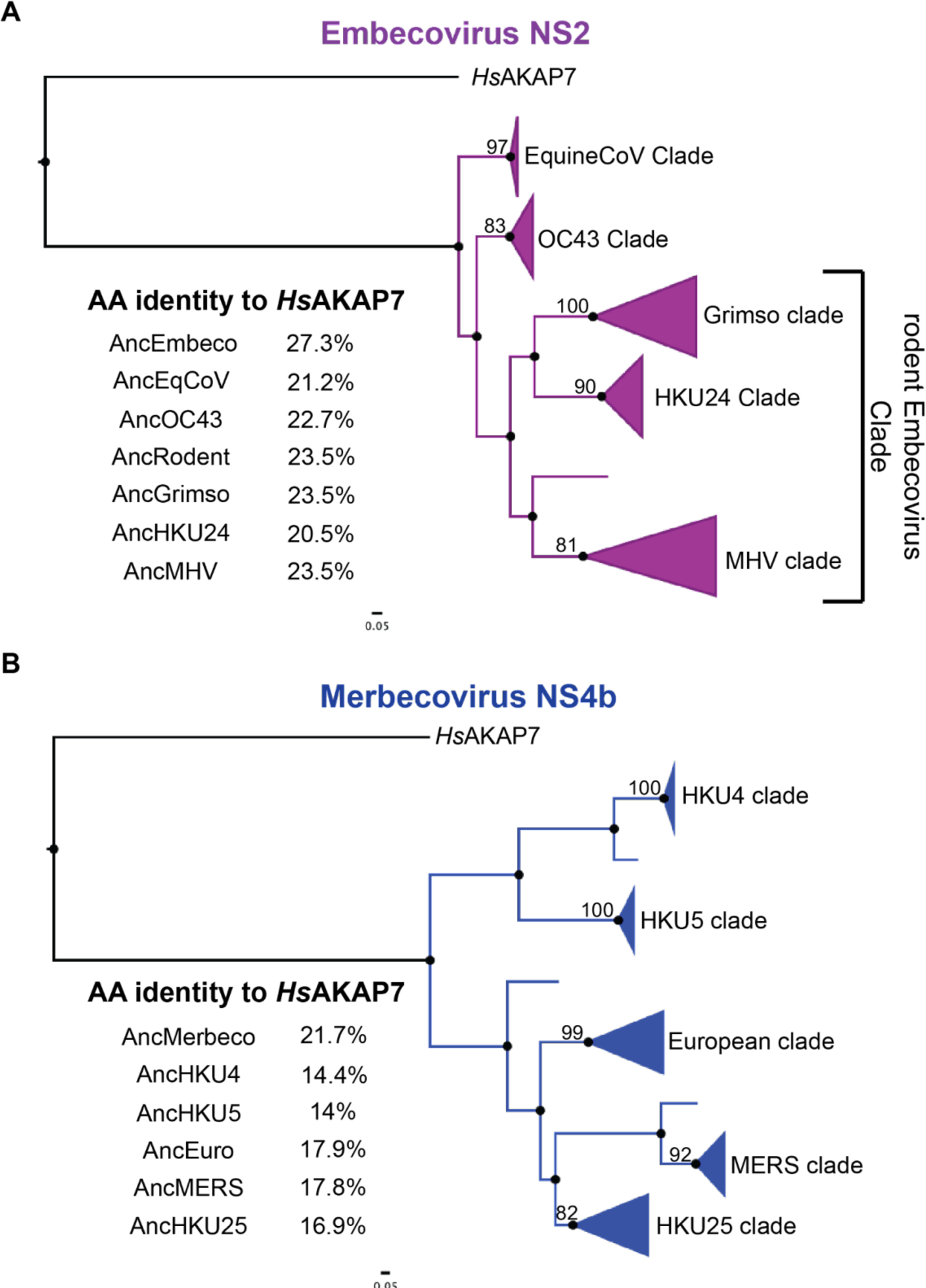
Ancestral sequence reconstruction (ASR) of Embecovirus and Merbecovirus PDEs. A) ML tree of 55 Embecovirus PDE sequences, collapsed into major clades, rooted with the *Hs*AKAP7 PDE domain sequence (AA 82-287) and sequence identity to *Hs*AKAP7 of reconstructed sequences at major nodes. B) ML tree of 27 Merbecovirus PDE sequences, collapsed into major clades, rooted with the *Hs*AKAP7 PDE domain sequence (AA 82-287) and sequence identity to *Hs*AKAP7 of reconstructed sequences at major nodes.

We conducted the same analysis with 28 Merbecovirus NS4b sequences, which was of particular interest due to the wide range in size of these proteins (246-285 AA), largely due to a highly variable N-terminus. Ancestral sequence reconstruction produced an AncMerbecovirus (AncMerbeco) NS4b with 21.7% identity to *Hs*AKAP7, again substantially higher than for ancestral sequences at reconstructed nodes or among observed sequences (**Figure 3B**). The AncMerbeco NS4b is 246 AA long, the same as human MERS-CoV isolates and significantly shorter than the 285 AA NS4b encoded by most HKU4 isolatesWe also reconstructed ancestral sequences of Alphacoronavirus and Torovirus PDEs. In the case of Alphacoronaviruses, this allowed comparisons not only with *Hs*AKAP7 but also with ancestral Embecovirus PDEs, critical for defining the relationship between them (**Figure S2**).

### Nidovirus PDE structural models show similar divergence from *Hs*AKAP7

Using proposed ancestral sequences, we generated Nidovirus PDE structural models and compared them to the structure of human (*Hs*)AKAP7 PDE to determine if there were differences in root mean square deviation (RMSD), a quantitative estimate of the similarity of structures. Overall amino acid identity was the same between AncEmbeco (**Figure 4A**) and AncAlphaCoV (**Figure 4B**) compared to *Hs*AKAP7 (∼27%) and RMSDs relative to the *Hs*AKAP7 structure were nearly identical, indicating similar divergence from a cellular ancestor and suggesting they may have a common origin, supporting a scenario of horizontal gene transfer between these clades. In contrast, AncMerbeco NS4b has the lowest amino acid identity to *Hs*AKAP7(∼21%) and highest RMSD (**Figure 4C**), supporting its independent origin which, given the substantial within-clade diversity of these proteins (**Figure 2A**), indicates an ancient acquisition event. In contrast, the torovirus PDE encoded on the 3’ end of Orf1a has relatively high identity to *Hs*AKAP PDE (∼39%) and the lowest RMSD value (**Figure 4D**). Coupled with the limited within-clade diversity among torovirus PDEs (**Figure 2A**) this suggests the torovirus PDE was acquired independently and more recently than the other Nidovirus PDEs.

**Figure 4.**
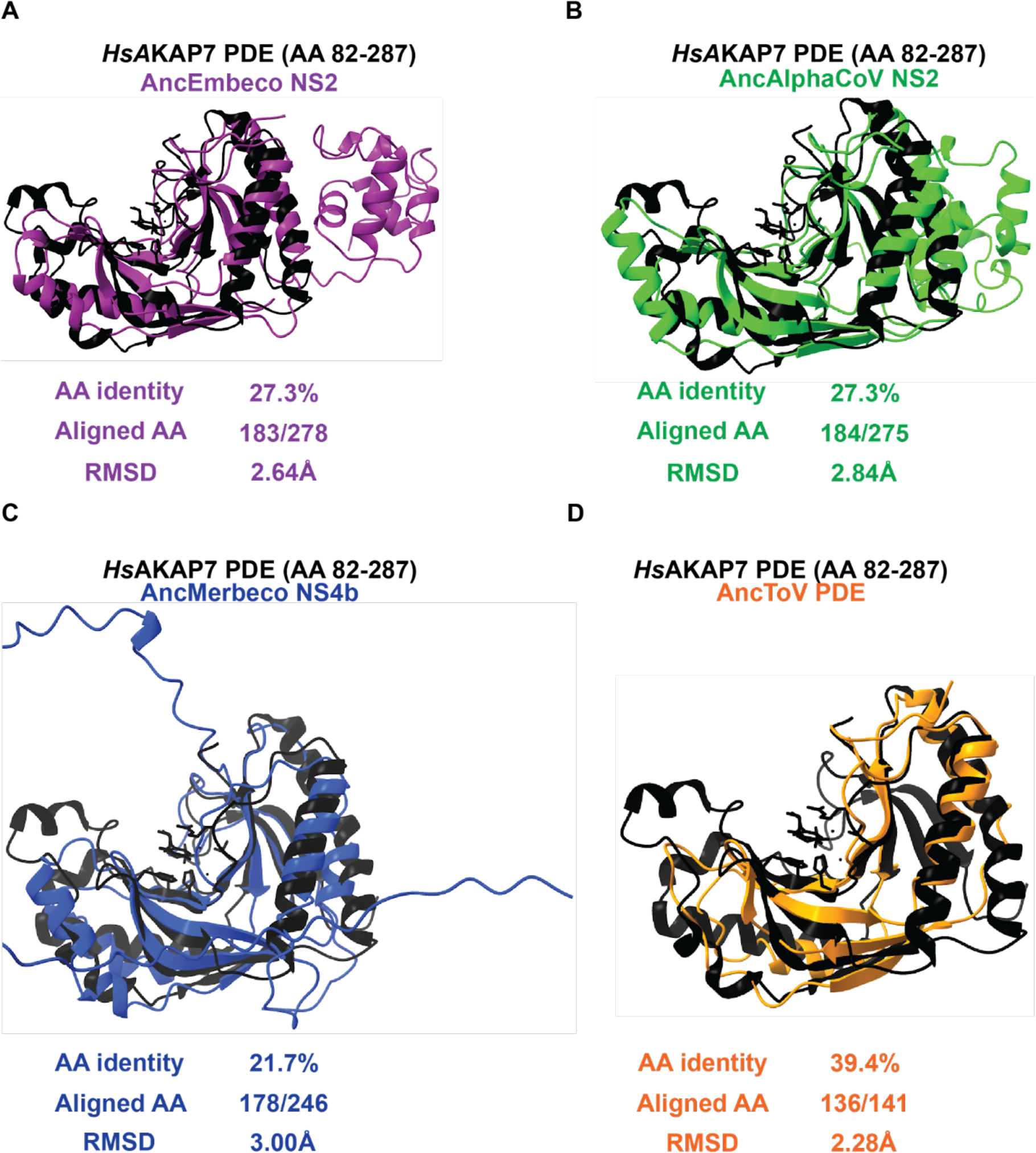
Structural similarity between *Hs*AKAP7 and Nidovirus PDEs. A) Overlay of the AncEmbecovirus PDE structure predicted in AlphaFold2 with the solved structure of the *Hs*AKAP PDE. The number of aligned amino acids (from the query sequence) and the RMSD were calculated using FatCat. B) Overlay of the AncAlphacoronavirus PDE structure predicted in AlphaFold with the solved structure of the *Hs*AKAP PDE. The number of aligned amino acids (from the query sequence) and the RMSD were calculated using FatCat. C) Overlay of the AncMerbecovirus PDE structure predicted in AlphaFold with the solved structure of the *Hs*AKAP PDE. The number of aligned amino acids (from the query sequence) and the RMSD were calculated using FatCat. D) Overlay of the AncTorovirus PDE structure predicted in AlphaFold with the solved structure of the *Hs*AKAP PDE. The number of aligned amino acids (from the query sequence) and the RMSD were calculated using FatCat.

A potentially complicating feature of the analysis is insertions and deletion events in PDEs. Relative to the *Hs*AKAP7 PDE structure, which comprises just the central 206 AA of the 315 AA *Hs*AKAP7 protein, Nidovirus PDEs have additional domains at the N-termini (Merbeco) or C-termini (Embeco and AlphaCoV). To account for the possibility that RMSD values might be inflated by poor modeling of these domains, we predicted the structure of the core PDE domains of Nidovirus PDEs. The AncMerbeco core PDE was not successfully predicted by AlphaFold2, so we used the sequence from a more recent node, AncMERS PDE. Overall, similarity to *Hs*AKAP7 PDE was generally unchanged regardless of whether the full-length or core viral PDE structural model was analyzed (**Figure S3**), bolstering the value of comparing full-length PDE structural models despite variable and disordered termini.

### Motif and structural analysis support Embecovirus-to-Alphacoronavirus horizontal transfer of NS2

Our prior phylogenetic and structural analyses were consistent with a relationship between Embecovirus and Alphavirus PDEs, such that one was derived from the other via recombination. Because these viruses fall into different coronavirus genera, a single introduction event is implausible, leaving the possibility of a recombination event or acquisition of a similar cellular ancestor. Given the longer branch lengths in the Embecovirus phylogeny, we hypothesize the acquisition event occurred in ancestral Embecoviruses and that Orf2 was subsequently transferred to the Alphacoronavirus clade. To refine our understanding of this history more deeply, we conducted additional sequence and structural analysis.

We first analyzed conservation of key motifs between AncEmbeco NS2 and the reconstructed ancestral PDE sequences from the other viral clades. Compared to AncEmbeco NS2, AncAlphaCoV NS2 had the fewest amino acid differences, with only three substitutions outside the catalytic His motif in which the second residue frequently toggles among PDEs (**Figure 5A**), supporting the phylogenetic inference that the Embecovirus and Alphacoronavirus NS2 proteins are closely related. We then conducted a series of overlays of AlphaFold2-predicted structural models of reconstructed PDEs as well as the solved *Hs*AKAP7 PDE structure. Specifically, we used a core PDE domain structural model to capture similarities/differences in the PDE fold itself and avoid disordered regions or N-terminal/C-terminal domains that AlphaFold2 failed to model. Modeling the core region better approximates the *Hs*AKAP7 PDE structure which was solved for just the 206 AA PDE central domain, not the entire protein.

**Figure 5.**
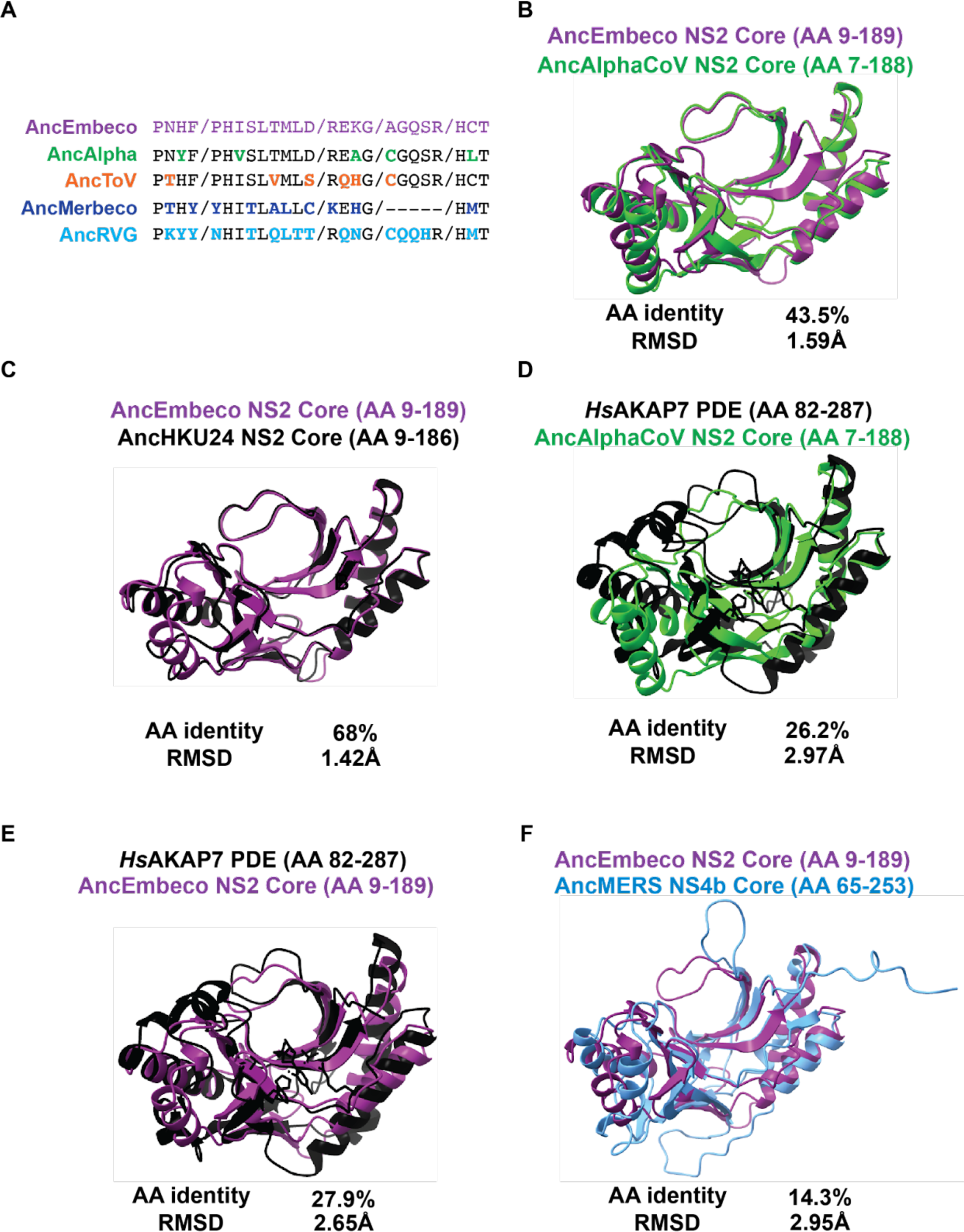
Comparative analysis of AncEmbeco and AncAlphaCoV PDEs. A) Motif analysis with AncEmbeco as the reference. Amino acid differences in key motifs are bolded and in the color corresponding to the viral PDE. B) Overlay and RMSD comparison of AncEmbeco NS2 and AncAlphaCoV NS2. C) Overlay and RMSD comparison of AncEmbeco NS2 and AncHKU24 NS2. D) Overlay and RMSD comparison of *Hs*AKAP PDE and AncAlphaCoV NS2. E) Overlay and RMSD comparison of *Hs*AKAP PDE and AncEmbeco NS2. F) Overlay and RMSD comparison of AncEmbeco NS2 and AncMERS NS4b.

The RMSD recovered by comparing the AncEmbeco and AncAlphaCoV models was comparable to the RMSD of AncEmbeco and AncHKU24, from a more recent inferred node in the Embecovirus PDE phylogeny (**Figure 5B, 5C**) although amino acid identity was substantially lower. Comparing the AncEmbeco and AncAlphaCoV NS2 models to the *Hs*AKAP PDE structure revealed that the viral PDEs are much closer to each other in amino acid identity and RMSD than either is to *Hs*AKAP7 PDE (**Figure 5D, 5E)**. Although it’s possible this is due to convergence in their structural evolution, the combination of higher amino acid identity and structural model similarity strongly support a common origin, which resulted from recombination. Finally, we compared the AncEmbeco NS2 model to the AncMERS NS4b model (AlphaFold2 failed to predict the AncMerbeco core PDE structure), showing that their divergence is comparable to that between the NS2 PDEs and *Hs*AKAP7 PDE (**Figure 5F**) and supporting a single origin of the embecovirus and alphacoronavirus PDEs.

### Merbecovirus PDEs exhibit significant intra-clade diversification

In addition to untangling the origin of viral PDEs, we sought to understand how they evolved after acquisition as new virus-encoded proteins compared to their eukaryotic ancestors. Merbecovirus NS4b stands out among other viral PDEs for its striking intra-clade diversification and structural divergence. In contrast, Embecoviruses exhibit shorter branch lengths (**Figure 2B**) and have higher sequence identity, even when accounting for greater sampling in the dataset and supporting their use for comparisons with Merbecovirus NS4b. Relative to the AncEmbeco NS2, two reconstructed sequences at recently inferred nodes have high sequence conservation (**Figure 6A**) and the modeled structures have a low RMSD value, suggesting minimal structural change throughout the course of Embecovirus NS2 evolution (**Figure 6B**). We also compared how the radiation of Embecovirus NS2 altered structure across the phylogeny by overlaying the modeled structures of the reconstructed AncMHV and AncGrimso NS2 proteins (**Figure 6C**). This confirmed that PDE structural divergence is relatively constrained even among diverse Embecoviruses.

**Figure 6.**
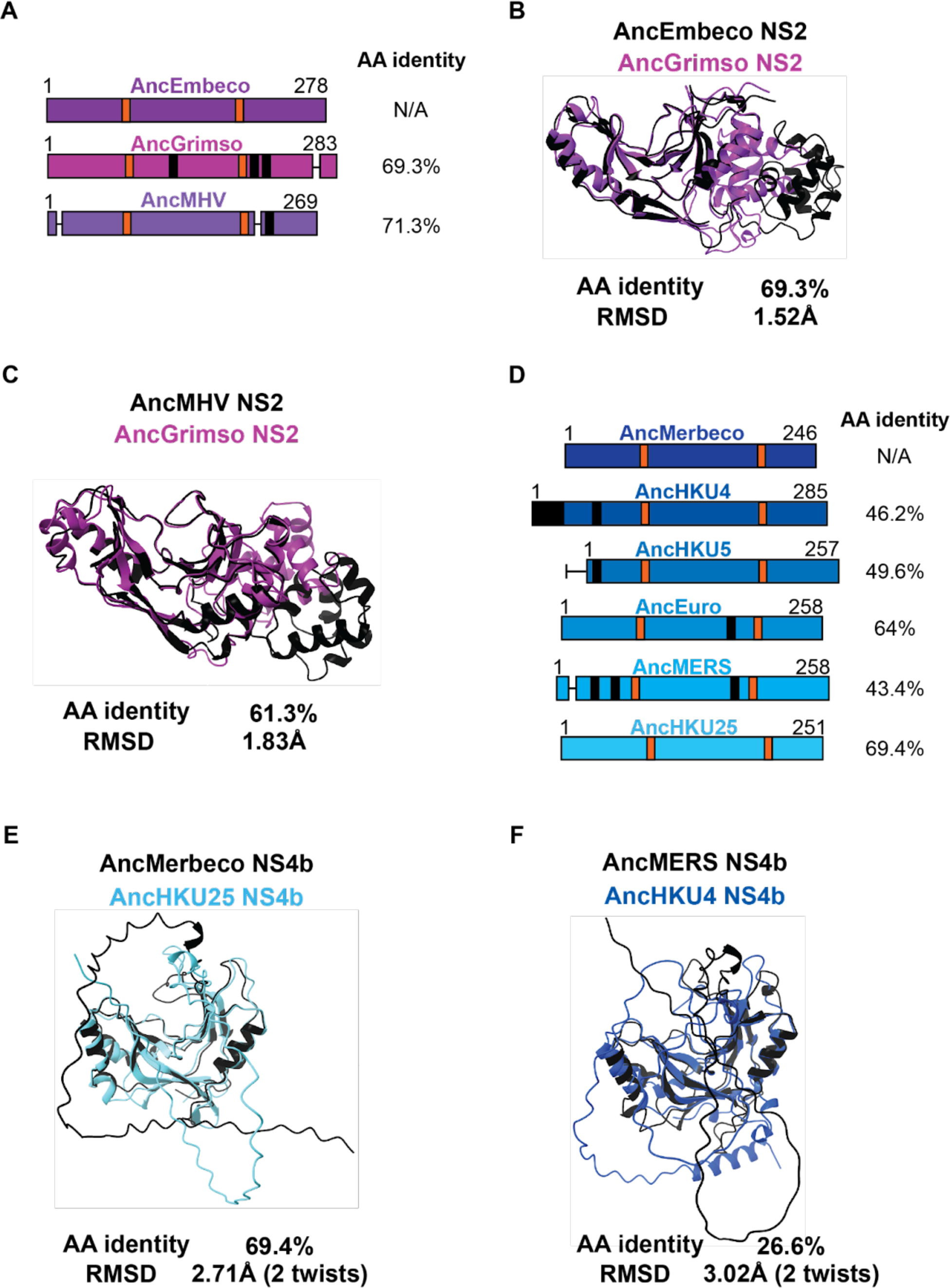
Variable sequence and structural PDE diversity within Nidovirus subgenera. A) Schematic of Embecovirus NS2 at key ancestral nodes with catalytic motifs (orange), deletions (horizontal lines), and insertions (black) indicated along with protein length and amino acid identity to the AncEmbeco sequence. B) Overlay of AlphaFold predicted structures of the reconstructed AncEmbeco and derived AncGrimso sequences with amino acid identity and the RMSD as calculated in FatCat. C) Overlay of AlphaFold predicted structures of the AncMHV and AncGrimso sequences with amino acid identity and the RMSD as calculated in FatCat. D) Schematic of Merbecovirus NS4b at key ancestral nodes with catalytic motifs (orange), deletions (horizontal lines), and insertions (black) indicated along with protein length and amino acid identity to the AncMerbeco sequence. E) Overlay of AlphaFold predicted structures of the AncMerbeco sequence and a shallower node (AncHKU25) with similar AA identity to each other as between the sequences in panel B, with RMSD from a FatCat comparison. F) Overlay of AlphaFold predicted structures of shallower ancestral nodes with different AA sequence identity and RMSDs from FatCat comparisons.

In contrast, the Merbecovirus NS4b has undergone significantly greater sequence divergence and structural variation. There has been notable extension, truncation, insertion, and deletion in the N-terminus of the protein (**Figure 6D**), likely facilitated by the fact that NS4b is encoded on a bicistronic ORF4ab viral sgRNA, which enables it to tolerate mutations that move the start codon upstream or downstream of the ancestral site within the overlapping region with Orf4a. However, even accounting for sequence divergence, the structural variation among Merbecovirus NS4b proteins is remarkable. AncMerbeco and AncHKU25 NS4b (**Figure 6E**); reconstructed at a less basal node) exhibit 69.4% amino acid identity, comparable to AncEmbeco and AncGrimso (**Figure 6B**) yet their RMSD is 2.71Å compared to 1.52Å for the Embecovirus NS2 proteins and the deviation of reconstructed sequences at recently inferred nodes of the Embecovirus phylogeny is still 1.83Å (**Figure 6C**). The reconstructed sequences of AncMERS and AncHKU4 NS4b exhibit just 26.6% amino acid identity and an average distance of deviation >3Å, reflecting marked diversification among Merbecovirus PDEs (**Figure 6F**).

### Sequence and structural analysis of Rotavirus PDEs supports two horizontal transfer events

Rotavirus PDE and VP3 phylogenies are consistent with independent acquisitions of PDEs in the RVA VP3 ancestor and in a common ancestor of RVB and RVG VP3 (**Figure 2A**, 2C). As with the nidovirus PDEs, ancestral sequence reconstruction was successful in producing sequences with the highest amino acid identity to *Hs*AKAP7 at the deepest node of the tree (**Figure 7A, 7C**). The AncRVA and AncRVB/G PDEs exhibited similar amino acid identity and RMSDs to *Hs*AKAP7 of 34-36% and ∼2.4-2.8Å (**Figure** 7B, 7D). However, their amino acid identity to each other was considerably lower than to *Hs*AKAP7 PDE and their modeled structures were also dissimilar, with RMSD >3Å (**Figure 7E**). This result strongly suggests that these PDEs are similarly diverged from *Hs*AKAP7 and each other, consistent with multiple origins of the rotavirus PDEs despite their synteny. Comparison of modeled structures of reconstructed RVB and RVG sequences reveal a stark contrast (**Figure 7F**). These reconstructed PDE sequences had nearly 50% amino acid identity and an RMSD <1Å, consistent with the phylogenetic inference that the RVB and RVG PDE arose from a single horizontal transfer event in a common ancestral VP3.

**Figure 7.**
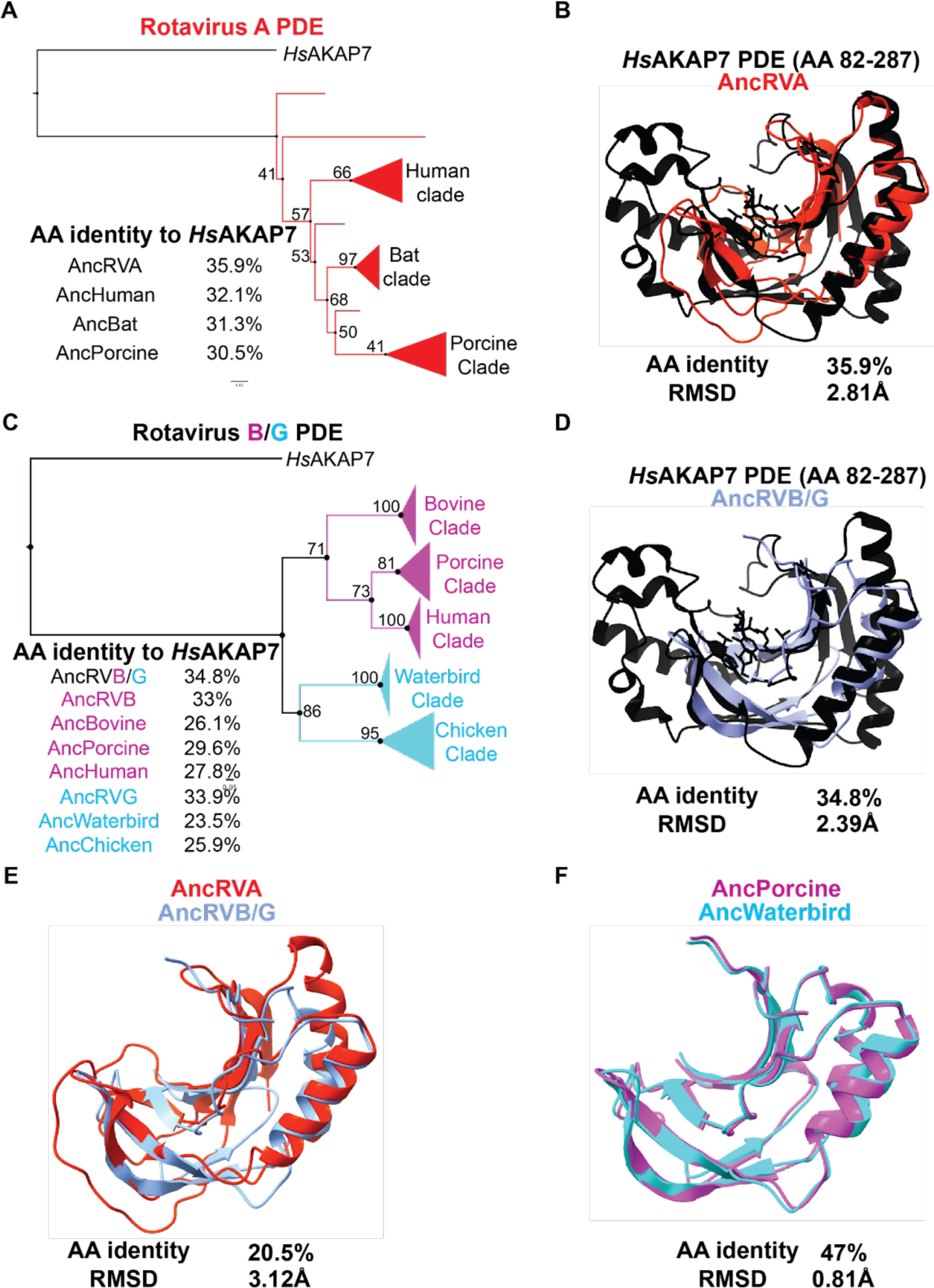
Ancestral sequence reconstruction and structural analysis of Rotavirus PDEs. A) Ancestral sequence reconstruction (ASR) of RVA PDEs at major nodes with AA identity to *Hs*AKAP B) Overlay of the AncRVA PDE structure predicted in AlphaFold with the solved structure of the *Hs*AKAP PDE. C) ASR of RVB/G PDEs at major nodes with AA identities to the hAKAP7 PDE sequence. D) Overlay of the AncRVB/G PDE structure predicted in AlphaFold with the solved structure of the *Hs*AKAP PDE. E) Overlay of the AncRVA PDE structure modeled in AlphaFold with the structural model of the AncRVB/G PDE. E) Overlay of the AncRVB Porcine clade PDE model structure and AncRVG Waterbird clade PDE structural model.

### Loss of PDE function in Rotavirus G is not correlated with major structural changes

A previous study compared enzymatic activity of PDEs encoded by RVA, RVB, and RVG and found that the PDE of the RVG tested was inactive (11). Although aT90N amino acid substitution in the second catalytic motif is associated with loss of enzymatic activity, reversion of the mutation did not rescue function. These findings point to cryptic functional determinants outside the catalytic sites, as previously reported for MHV NS2 (18). The PDE sequences from AncRVG and AncChicken RVG contain identical key motifs and threonines at position 90 (T90; **Figure 8A**). In contrast, RVG^inactive^ and other closely related viruses encode substitutions at position 90 and in other conserved motifs (**Figure 8A**). To determine whether structural divergence might underlie loss-of-function we compared structural models of RVB^active^ and RVG^inactive^, which are similarly diverged from the AncRVB/G model in both sequence identity and RMSD (**Figure** 8B, 8C). We found that RVB^active^ and RVG^inactive^ are similar to each other (**Figure 8D**) and therefore no obvious structural changes can explain the loss of function in the PDE of RVG. Whether amino acid changes in the key conserved motifs are involved is unclear and an important topic for future experimental study.

**Figure 8.**
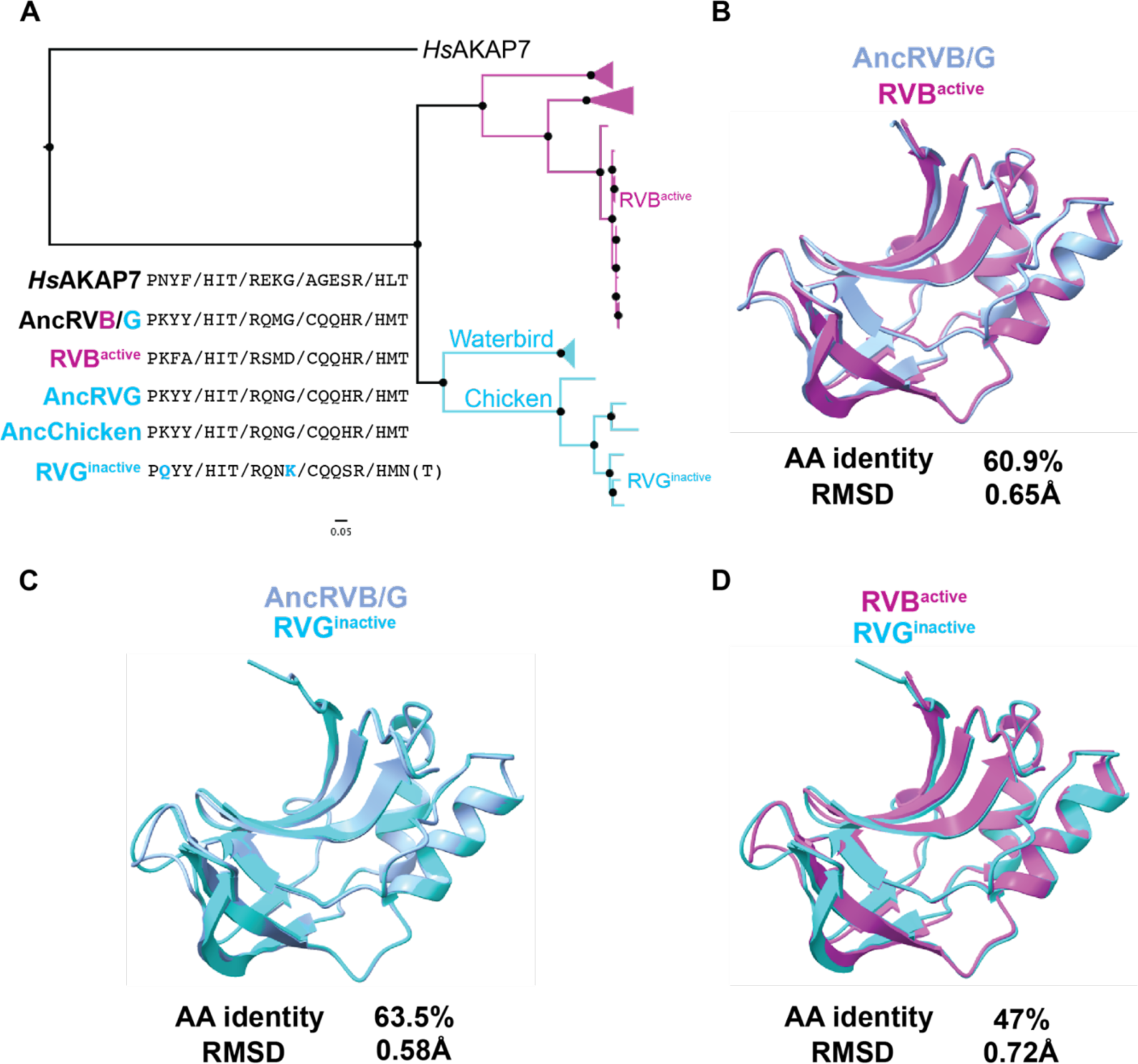
Sequence motif and structural analysis of RVB and RVG functional divergence. A) ML tree of RVB and RVG PDEs, with previously assayed PDEs (Ogden et. al. 2015) indicated B) Overlay and similarity metrics for the predicted structures of the AncRVB/G PDE and the active RVB PDE tested by Ogden et. al. C) Overlay and similarity metrics for the predicted structures of the AncRVB/G PDE and the inactive RVG PDE tested by Ogden et. al. D) Overlay and similarity metrics for the predicted structures of the active RVB PDE and the inactive RVG PDE tested by Ogden et. al. RMSDs for all overlays were calculated in FatCat.

## Discussion

In this study we traced and characterized the acquisition and evolution of viral phosphodiesterases. These proteins are among the clearest examples of acquisition of a host gene by RNA viruses, in contrast to more abundant examples among DNA viruses. Previous work examining the role of gene flow in RNA virus evolution has largely considered conserved genes such as capsids, RNA-dependent RNA polymerases, and helicases (19). In contrast, we characterize an example of gene transfer shaping virus interactions with host immunity, with potential implications for host-range and virulence. A recent report proposing bacterial origins for eukaryotic OAS genes, suggests that both sides of the interface between viral PDE and OAS-RNase L arose via horizontal gene transfer (20). Understanding how HGT has influenced virus-innate immune interfaces benefits from a better understanding of the emergence of newly captured viral genes and their subsequent evolution. Here, we took advantage of recent advances in protein structure prediction to study the evolution of viral PDEs.

The amino acid sequence of viral PDEs can undergo extensive substitution without loss of function so identity between viral and host sequences is often very low. This complicates phylogeny-only approaches to understanding the history of these proteins, which motivated our integration of sequence and structural prediction-based comparisons. Previous work has linked these proteins to an AKAP7-like ancestor (2,16) based on similarity to solved *Hs*AKAP7, MHV NS2, and RVA PDE structures (9–11,21) but these analyses provide only a limited view of PDE evolution.

Our phylogenetic analysis is a major expansion on prior efforts (16), incorporating 173 PDE sequences from Embecoviruses, Merbecoviruses, Toroviruses, Alphacoronaviruses, RVAs, RVBs, and RVGs. The arrangement of the PDE tree, including several informative long branches in comparison to the RdRp tree, suggests multiple independent horizontal transfer events and one virus-to-virus transfer via recombination, accounting for incongruence with the Nidovirus RdRp tree (**Figure 2B**). Comparison of the rotavirus PDE and VP3 phylogenies supports two introductions, one into an ancestral RVA VP3 and one into an RVB/G VP3 common ancestor (**Figure 2C**). The syntenic but independent acquisition of PDEs by Rotaviruses suggests that the VP3 segment is particularly tolerant of gaining new genetic material. Future studies could test this idea by experimentally approximating horizontal transfer events using the established rotavirus A molecular clone (8,22).

Our characterization of several horizontal transfer events of orthologous ancestral AKAPs and their common use in antagonism of the OAS-RNase L pathway has intriguing biological implications. The cellular role of the AKAP7 PDE remains mysterious, so whether these proteins were evolutionarily repurposed after viral acquisition is currently unclear. Importantly, these horizontal transfer events are recurrent but phylogenetically restricted, occurring multiple times only in Rotaviruses and Nidoviruses despite the fact that the OAS-RNase L pathway restricts diverse viral families (24). Among RNA viruses, constraint on genome size due to error rate and packaging capacity may be a limiting factor (25), which is alleviated among coronaviruses by their proofreading exonuclease and resultant large genomes.

The ability of some coronaviruses to tolerate insertions of PDE genes raises the question of why other coronavirus subgenera, including sarbecoviruses such as SARS-CoV-2, lack PDEs. Two possibilities can be proposed. One is that OAS-RNase L activity, and thus the selective pressure to encode an antagonist is cell-type specific (12,26). The cellular tropism of sarbecoviruses in their natural rhinolophid hosts is unknown, but it is possible they primarily infect cells that lack a potent OAS-RNase L pathway, similar to MHV and liver hepatocytes (12). Alternatively, rhinolophid bats may have acquired loss-of-function mutations in OAS genes, as seen with primate OAS1 (27), eliminating the pressure for sarbecoviruses to antagonize the pathway.

Merbecovirus NS4b highlights the potential for highly variable evolutionary trajectories following acquisition of hostPDE genes. They exhibit striking sequence divergence and structural deviation from both *Hs*AKAP7 and other viral PDEs and extensive divergence between each other. Relative to other viral PDEs and like AKAP7 proteins, Merbecovirus PDEs contain a nuclear localization sequence (NLS) and appear in the nucleus in abundance (7,28,29). However, the NLS is bipartite in contrast to the monopartite AKAP7 NLS and shows no evidence of being derived from the cellular ancestral NLS, suggesting itarose *de novo* early after acquisition of the PDE. Concordant with NS4b localization to the nucleus, this PDE has additional functions in immune antagonism distinct from disruption of RNase L activation (7,28–33). This multi-functionality distinguishes NS4b from all other viral PDEs and likely subjects NS4b to unique selective pressures, presumably accounting for its volatile evolutionary history.

As the evolutionary history of viral PDEs comes into better focus, our understanding of the roles for PDEs in viral replication and impacts on fitness remains limited. With the exception of MHV, any fitness benefits provided by coronavirus PDEs has not been explored, especially during infection and transmission of viruses in natural hosts. When bat coronaviruses move into new hosts the loss of some accessory protein function is common, suggesting the advantages provided are host specific. The development of more sophisticated cell culture systems from a variety of species, such as primary epithelial cells (34,35) and bat intestinal organoids (36) opens the possibility of unlocking new insights into how the combination of horizontal gene transfer and natural selection shapes the evolution of RNA viruses and their hosts.

## Methods

### Phylogenetic Analysis

#### Viral PDEs

All alignments (**Supplementary Materials**) were generated using the MAFFT plug-in (37) with default parameters in Geneious Prime v2022.2.1. Alignments were manually refined as necessary. Maximum-likelihood trees with 100 replicates were generated using IQ-Tree 2 (38) on the Los Alamos National Laboratory server. Different substitution models (with 20 rate-categories) were necessary for different alignments to maximize tree stability and branch support. The all-viral PDE tree is unrooted, as no clear outgroup exists to all viral PDEs. ML trees for individual clades of viral PDEs are rooted with *Hs*AKAP7. Alignments (.fasta*) and IQ-Tree logs (.log*) containing detailed parameters and associated with every PDE tree can be found in the Supplementary Files. The ViralPDEs.fasta file contains accessions for all viral PDE sequences analyzed.

#### Nidovirus RdRps

To generate the Nidovirus RdRp tree (**Figure 1B**) we aligned 145 Nidovirus RdRp sequences representing toroviruses, alphacoronaviruses, betacoronaviruses, gammacoronaviruses, and deltacoronaviruses using MAFFT (default parameters) and generated a ML tree in IQ-Tree 2 using a GTR20 substitution model, 20 rate categories, and 100 bootstrap replicates. Raw files associated with the analysis are in the Supplementary Files.

#### Rotavirus VP3

We aligned 137 rotavirus VP3 amino acid sequences using MAFFT and trimmed the PDE domains from the RVA, RVB, and RVG sequences in the alignment. We then used IQ-Tree 2 to generate a maximum-likelihood tree using a JTT substitution model, 20 rate categories, and 100 bootstrap replicates.

#### Maximum-likelihood tree visualization

All .treefile outputs from IQ-Tree 2 were imported into FigTree v1.4.4 for visualization and coloring. Branch support values and labels were added manually in Adobe Illustrator 2023.

### Sequence motif analysis

We used previously defined motifs (5,16) to identify motifs and guide manual scanning of the viral PDE/AKAP alignments.

### Ancestral Sequence Reconstruction and Analysis

We generated alignments of each viral PDE group with the *Hs*AKAP7 PDE domain (described above) and fed these alignments into FastML (39,40) along with the maximum-likelihood tree for each alignment generated with IQ-Tree 2. Ancestral sequences were then reconstructed using a JTT substitution model and other default parameters (optimize branch length, Gamma distribution, marginal reconstruction) and exported for analysis. The raw files (outFile_seq_marginal.txt*, outTreefileAncestor.txt*, outTreeFileNewick.txt*) are available on <u>FigShare</u>. After downloading the inferred ancestral sequences, we identified major nodes and aligned them to *Hs*AKAP7 to confirm that deeper nodes have higher amino acid percent identity to the eukaryotic reference PDE. We also analyzed evolutionary trajectory in conserved AKAP-like motifs.

### Structural Modeling

We exported major ancestral nodes from each viral PDE family as .fasta files and analyzed these using AlphaFold2 (v2.1.2) 9 as implemented at the University of Utah Center for High Performance Computing (CHPC) with the following parameters:

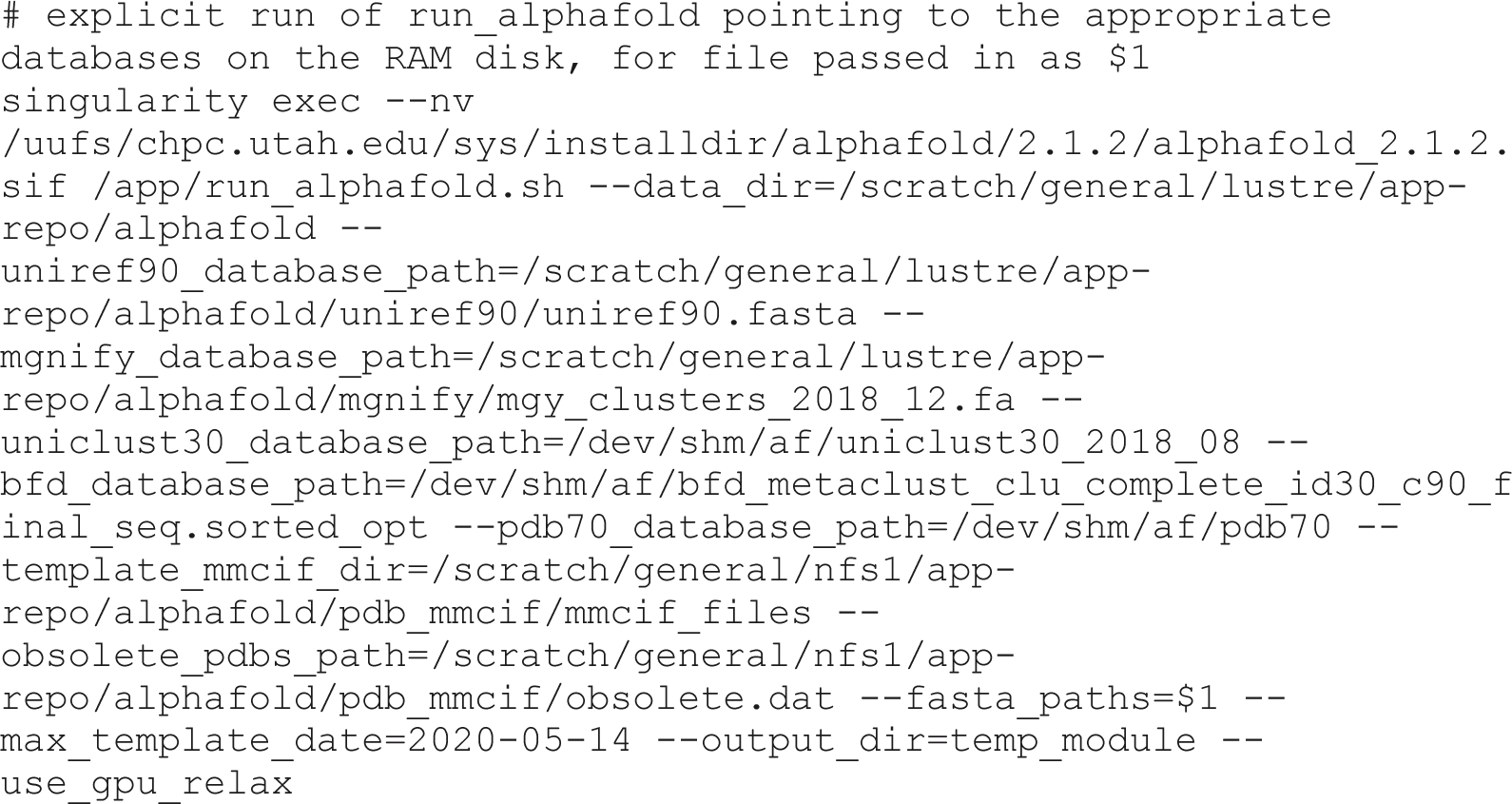

We then extracted the ranked_0.pdb model for each sequence for further analysis. Models were imported into and analyzed with UCSF ChimeraXv1.5 – developed by the Resource for Biocomputing, Visualization, and Informatics at the University of California, San Francisco, with support from National Institutes of Health R01-GM129325 and the Office of Cyber Infrastructure and Computational Biology, National Institute of Allergy and Infectious Diseases (41,42). Our primary analysis in ChimeraX involved overlays of predicted and solved structures to generate images that were then exported to Adobe Illustrator 2023.

To quantify the similarities (or differences) between different model structures we conducted pairwise alignments in FATCAT 2.0 (43) to calculate RMSD values.

## Figure Design

Figure 1 was produced using BioRender. All other figures were produced using Adobe Illustrator 2023.

## Acknowledgements

We thank Zoe A. Hilbert for advice and critical feedback. We thank Ian Boys for assistance implementing AlphaFold2 and FATCAT. We thank Kristina M. Babler for help designing the Figure 1 schematic. This work was supported by NIH grants F32AI152341 to S.A.G. and R35GM134936 to N.C.E.

---

**Figure S1.**
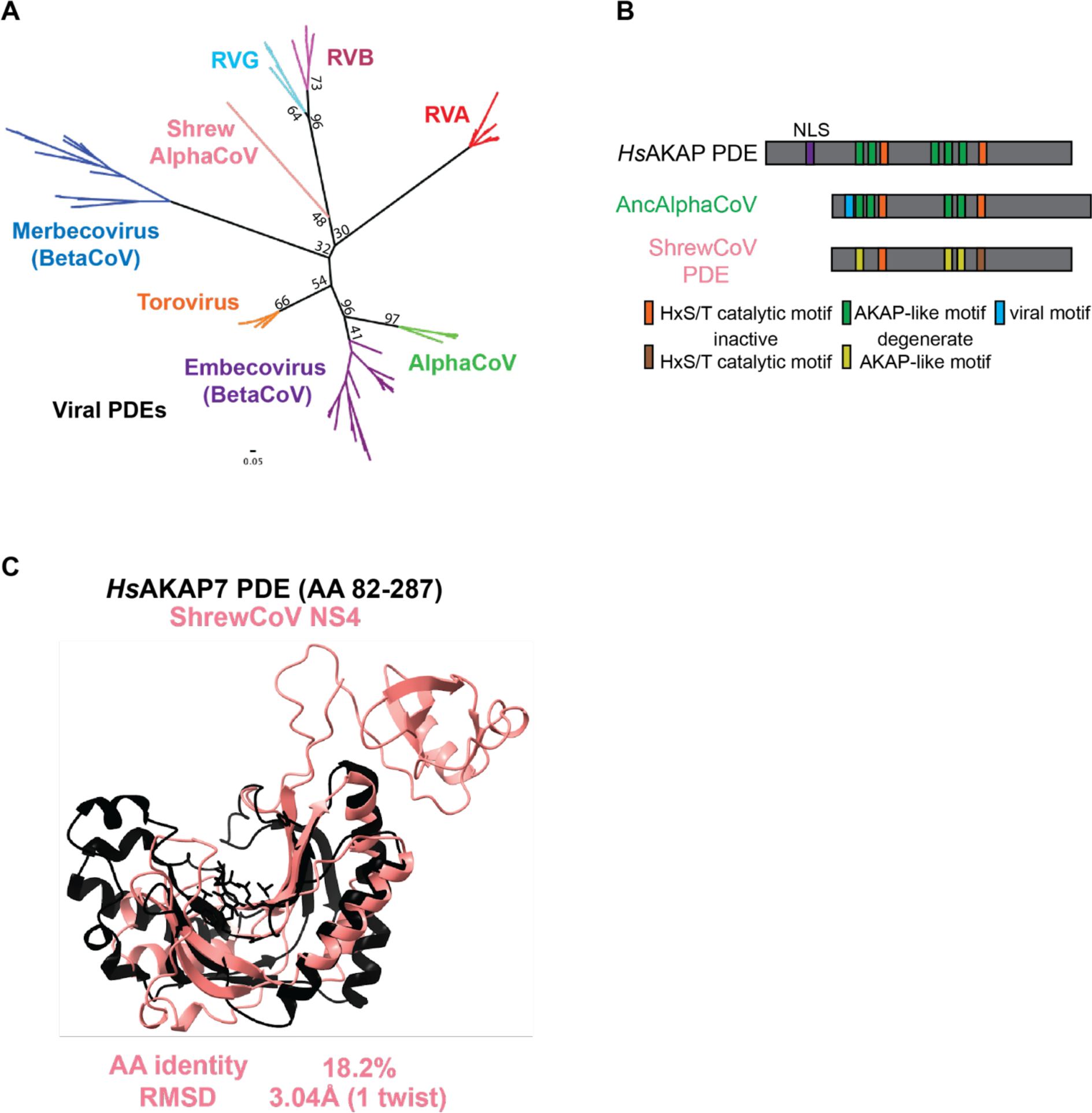
Phylogenetic and structural characterization of divergent, non-functional ShrewCoV PDE. A) ML tree of all virus PDEs, including the NS4 protein of ShrewCoV. B) Schematic of the ShrewCoV PDE in comparison to the *Hs*AKAP7 and rodent AlphaCoV PDEs, showing intact and degenerate but identifiable motifs. C) Overlay and RMSD comparison of *Hs*AKAP7 PDE and ShrewCoV NS4.

**Figure S2.**
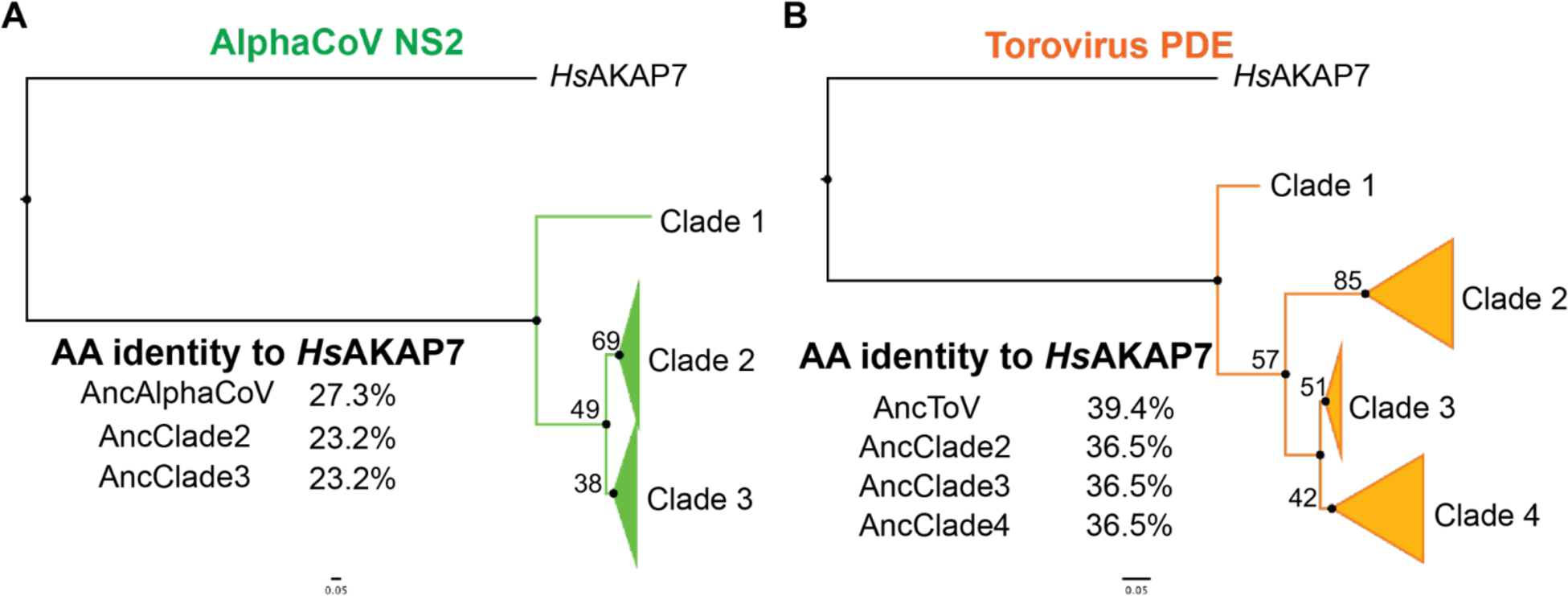
Ancestral sequence reconstruction (ASR) of Alphacoronavirus and Torovirus PDEs. A) ML tree of 10 Alphacoronavirus PDE sequences, collapsed into major clades, rooted with the *Hs*AKAP7 PDE domain sequence (AA 82-287) and sequence identity to *Hs*AKAP7 of reconstructed sequences at major nodes. B) ML tree of 18 Torovirus PDE sequences, collapsed into major clades, rooted with the *Hs*AKAP7 PDE domain sequence (AA 82-287) and sequence identity to *Hs*AKAP7 of reconstructed sequences at major nodes.

**Figure S3.**
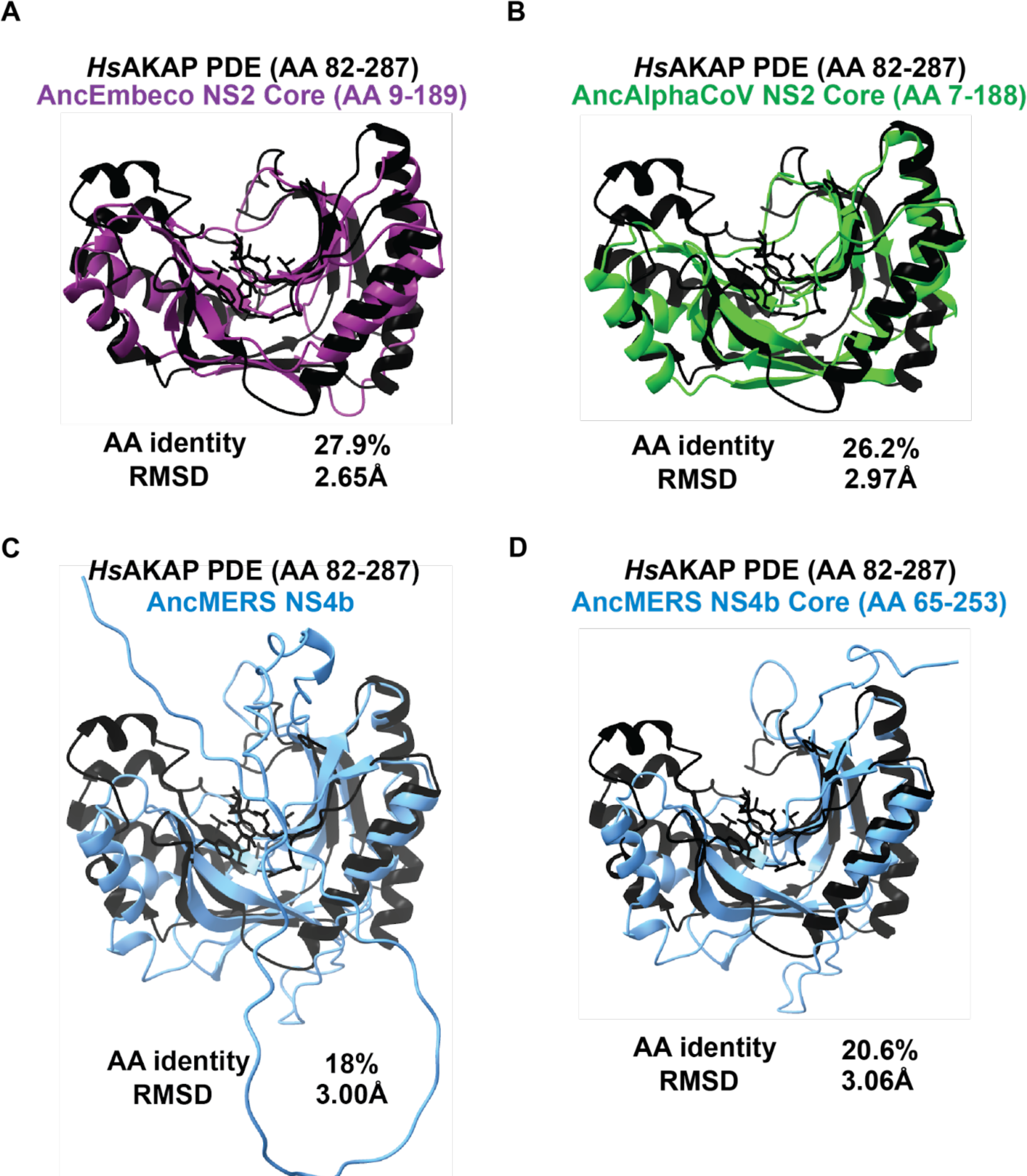
Sequence and structural divergence is not due to unique N and C termini of viral PDEs. A) Sequence and structural comparison of hAKAP PDE and the AncEmbeco NS2 core PDE predicted structure. B) Sequence and structural comparison of hAKAP PDE and the AncAlphaCoV NS2 core PDE predicted structure. C) Sequence and structural comparison of hAKAP PDE and the AncMERS NS4b PDE predicted structure. The AncMERS PDE was chosen because AlphaFold failed to predict a structure for the AncMerbeco core PDE domain D) Sequence and structural comparison of hAKAP PDE and the AncMERS NS4b core PDE predicted structure.

