## Supplemental Figures for "Recurrent Viral Capture of Cellular Phosphodiesterases that Antagonize OAS-RNase L"

A

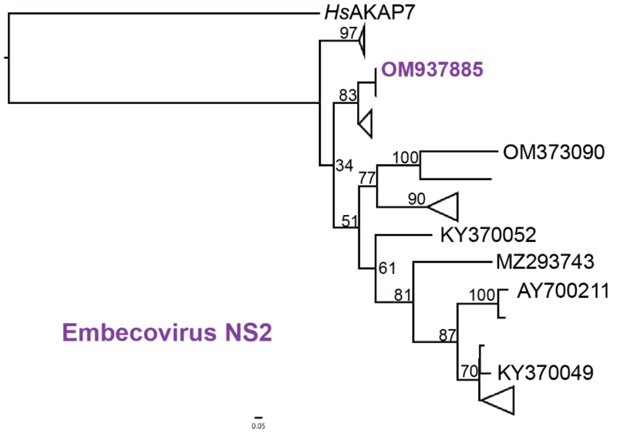


B

**Reference: OM937885**

| Virus | RdRp % identity | PDE % identity | Core PDE % identity |
|---|---|---|---|
| OM373090 | 86.3 | 47.2 | 47.6 |
| KY370052 | 94 | 61.1 | 60.8 |
| MZ293743 | 91.4 | 50.5 | 50 |
| AY700211 | 90.3 | 51.5 | 51.6 |
| KY370049 | 91.4 | 51.5 | 51.3 |

C

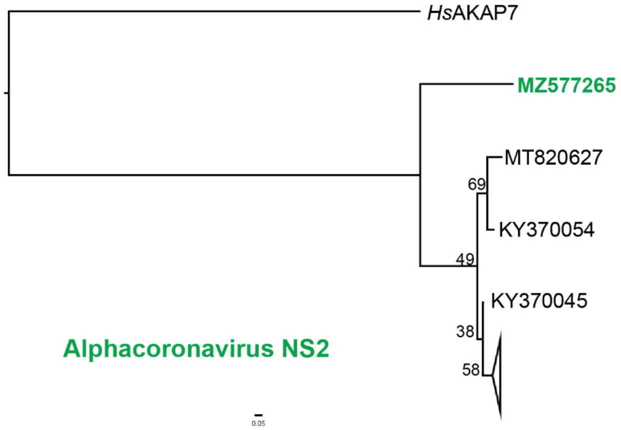


D

**Reference: MZ577265**

| Virus | RdRp % identity | PDE % identity | Core PDE % identity |
|---|---|---|---|
| MT820627 | 93.4 | 53.4 | 54.4 |
| KY370054 | 93.1 | 55.9 | 57.7 |
| KY370045 | 94 | 55.9 | 55.5 |

E

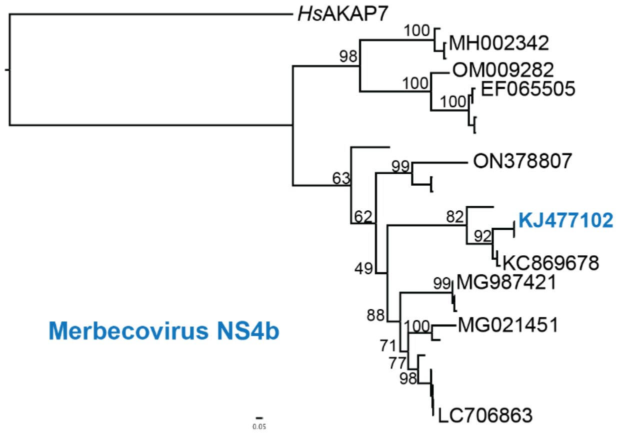


F

**Reference: KJ477102**

| Virus | RdRp % identity | PDE % identity | Core PDE % identity |
|---|---|---|---|
| MH002342 | 92.1 | 25.9 | 30.4 |
| OM009282 | 89.8 | 27.3 | 29 |
| EF065505 | 90 | 27 | 28.8 |
| ON378807 | 94.5 | 35.3 | 44.5 |
| KC869678 | 98.4 | 83.7 | 90.5 |
| MG987421 | 95 | 40.3 | 45.7 |
| MG021451 | 94.3 | 39.7 | 48.6 |
| LC786603 | 94.6 | 41.6 | 48.6 |

G

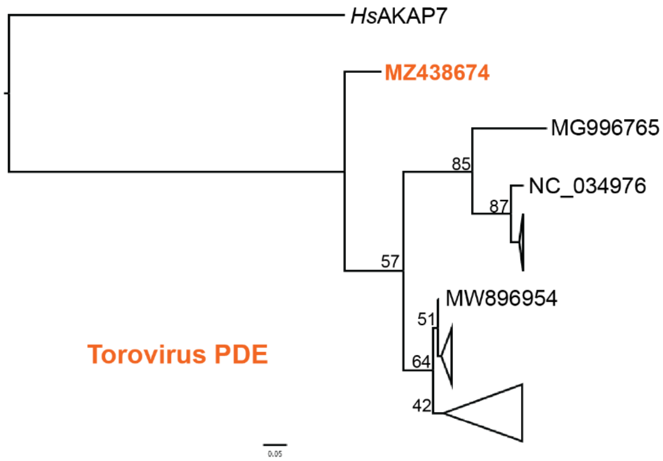


H

**Reference: MZ438674**

| Virus | RdRp % identity | PDE % identity |
|---|---|---|
| MG996765 | 83.5 | 71.6 |
| NC_034976 | 85.5 | 70.2 |
| MW896954 | 89.3 | 78.7 |

I

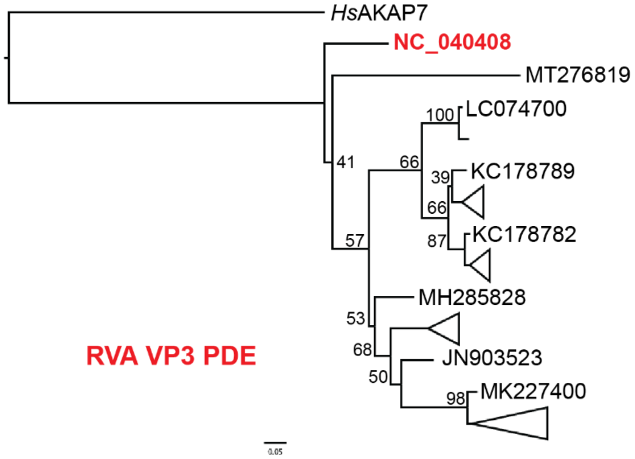


J

**Reference: NC_040408**

| Virus | VP3 % identity | PDE % identity |
|---|---|---|
| MT276819 | 71.6 | 66.9 |
| LC074700 | 74.6 | 73.2 |
| KC178789 | 74.1 | 72.5 |
| KC178782 | 75.4 | 71.1 |
| MH285828 | 76.8 | 77.5 |
| JN903523 | 75.8 | 72.5 |
| MK227400 | 76.5 | 73.2 |

K

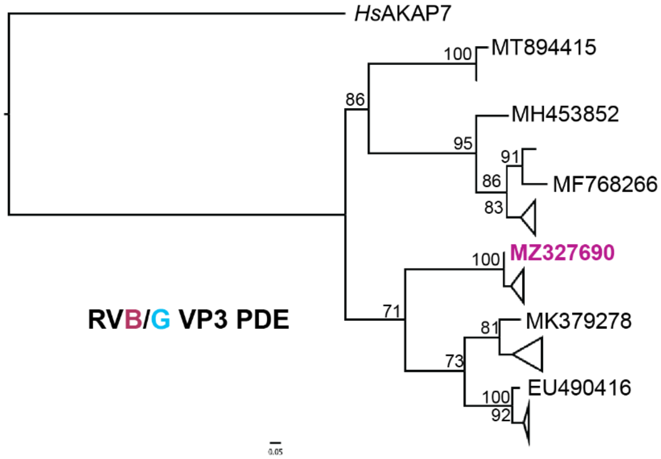


L

**Reference: MZ327690**

| Virus | VP3 % identity | PDE % identity |
|---|---|---|
| MT894415 | 49.7 | 50.4 |
| MH453852 | 44.1 | 44.3 |
| MF768266 | 46.1 | 47 |
| MK379278 | 62.5 | 53.9 |
| EU490416 | 63.6 | 54.8 |

**Figure S1. PDEs encoded by discrete open reading frames exhibit higher sequence divergence than sub-domain PDEs** A) ML tree of 55 Embecovirus NS2 AA sequences rooted with hAKAP7 PDE and with the reference sequence for AA identity comparisons colored and bolded B) Table of AA identity of select Embecovirus RdRp, PDE, and ‘Core’ PDE sequences to the reference sequence. C) ML tree of 10 Alphacoronavirus NS2 AA sequences rooted with hAKAP7 PDE and with the reference sequence for AA identity comparisons colored and bolded D) Table of AA identity of select Alphacoronavirus RdRp, PDE, and ‘Core’ PDE sequences to the reference sequence. E) ML tree of 27 Merbecovirus NS4b AA sequences rooted with hAKAP7 PDE and with the reference sequence for AA identity comparisons colored and bolded. F) Table of AA identity of select Merbecovirus RdRp, PDE, and ‘Core’ PDE sequences to the reference sequence. G) ML tree of 18 Torovirus PDE AA sequences rooted with hAKAP7 PDE and with the reference sequence for AA identity comparisons colored and bolded. H) Table of AA identity of select Torovirus RdRp, and PDE sequences to the reference sequence. I) ML tree of 35 Rotavirus A PDE rooted with hAKAP7 and with the reference sequence for AA identity comparisons colored and bolded. J) Table of AA identity of select Rotavirus A PDE and VP3 (minus the PDE) sequences to the reference sequence. K) ML tree of 20 Rotavirus B and 8 Rotavirus G PDE sequences rooted with hAKAP7 and with the reference sequence for AA identity comparisons colored and bolded. L) Table of AA identity of select Rotavirus B and G PDE and VP3 (minus the PDE) sequences to the reference sequence.

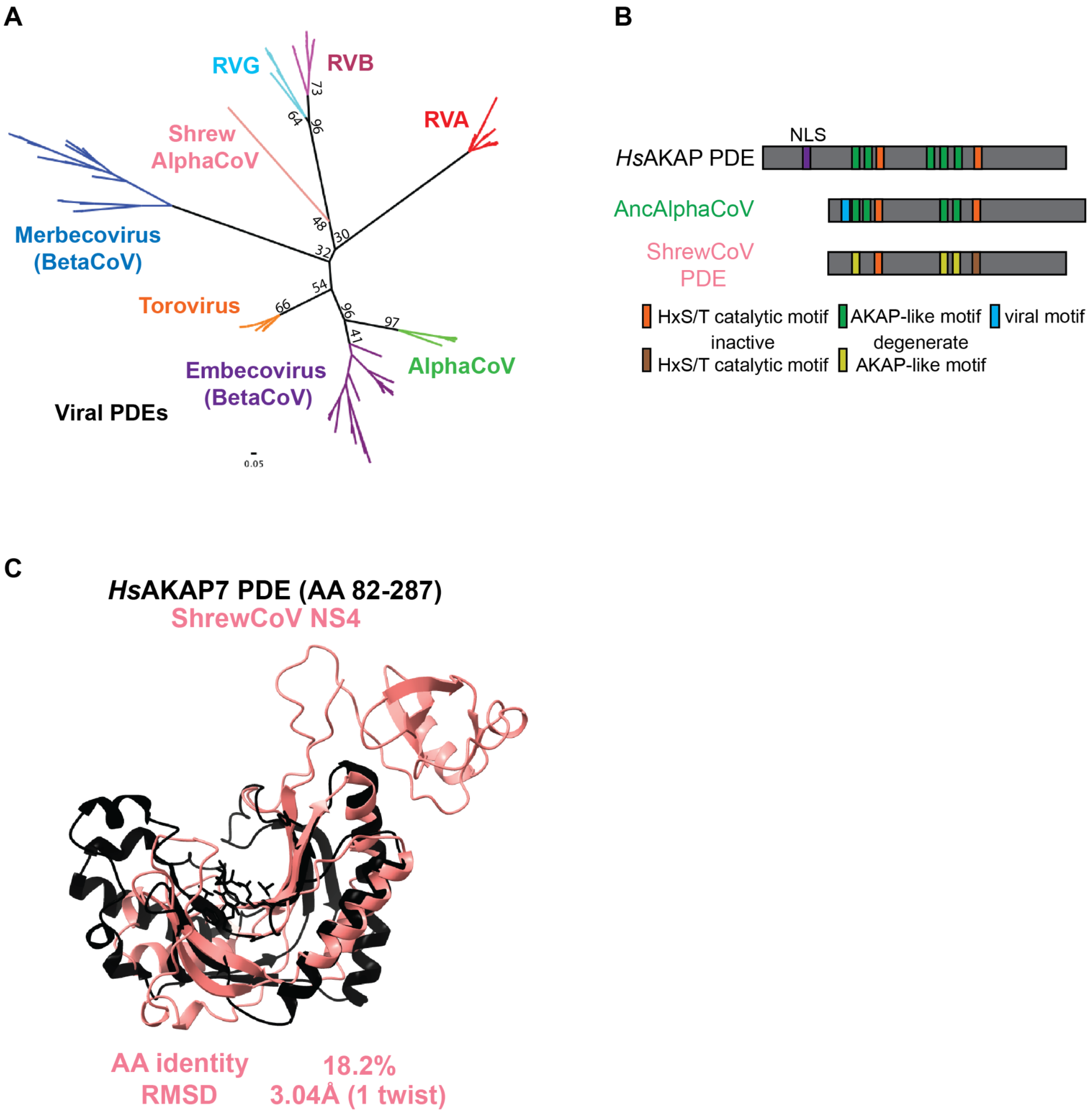


**Figure S3. Phylogenetic and structural characterization of divergent, non-functional ShrewCoV PDE.** A) ML tree of all virus PDEs, including the NS4 protein of ShrewCoV. B) Schematic of the ShrewCoV PDE in comparison to the *Hs*AKAP7 and rodent AlphaCoV PDEs, showing intact and degenerate but identifiable motifs. C) Overlay and RMSD comparison of *Hs*AKAP7 PDE and ShrewCoV NS4.

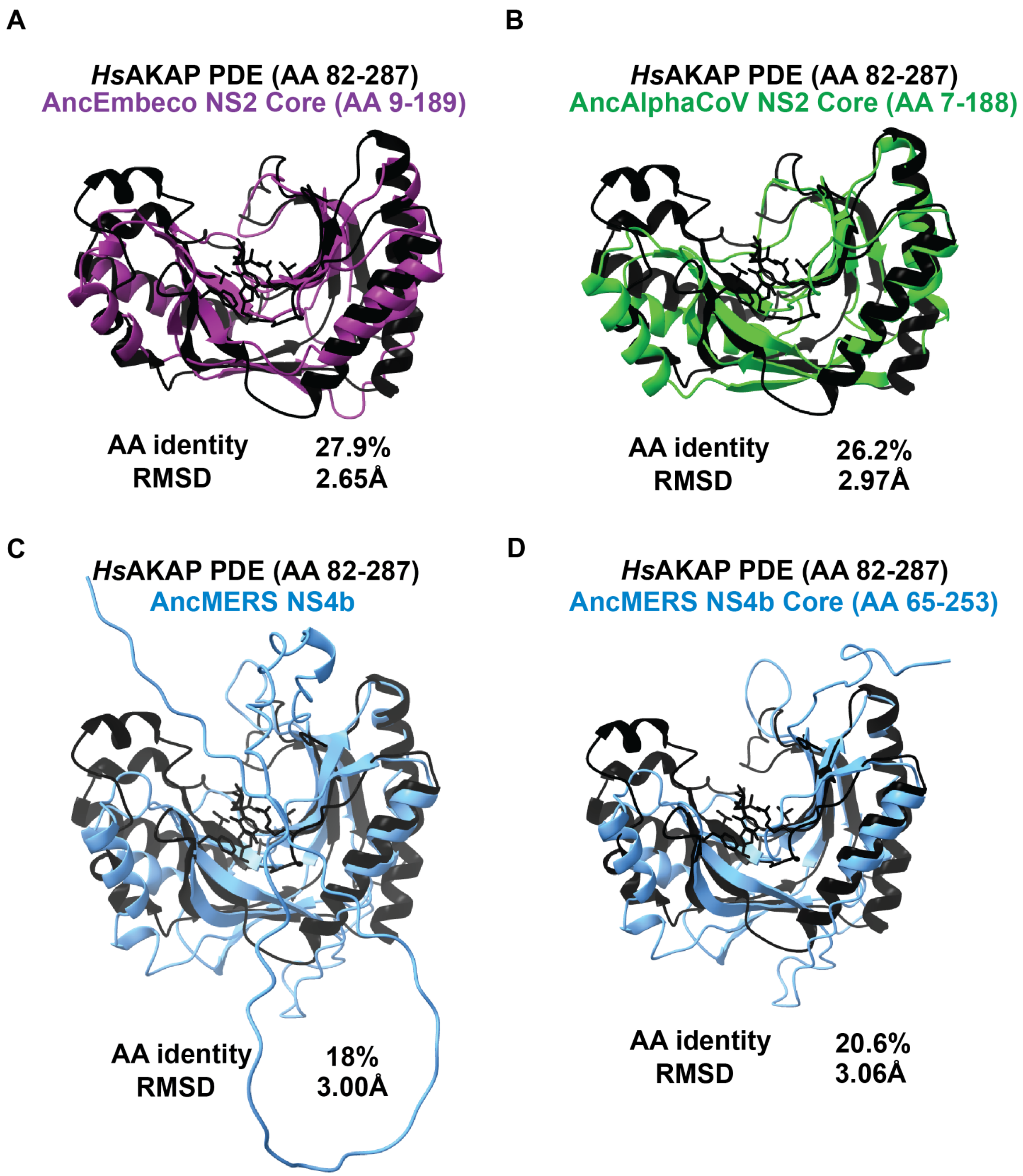


**Figure S4. Sequence and structural divergence is not due to unique N and C termini of viral PDEs.** A) Sequence and structural comparison of hAKAP PDE and the AncEmbeco NS2 core PDE predicted structure. B) Sequence and structural comparison of hAKAP PDE and the AncAlphaCoV NS2 core PDE predicted structure. C) Sequence and structural comparison of hAKAP PDE and the AncMERS NS4b PDE predicted structure. The AncMERS PDE was chosen because AlphaFold failed to predict a structure for the AncMerbeco core PDE domain D) Sequence and structural comparison of hAKAP PDE and the AncMERS NS4b core PDE predicted structure.
